# Loss-of-function of the drought-induced genes *GASA3* and *AFP1* confers enhanced drought tolerance in *Arabidopsis thaliana*

**DOI:** 10.1101/2025.04.03.647048

**Authors:** Sabarna Bhattacharyya, Bexultan Turysbek, Sebastian Lorenz, Diego Clavijo Rosales, Yasira Shoaib, Katharina Gutbrod, Peter Dörmann, Fatima Chigri, Ute C. Vothknecht

## Abstract

Prolonged drought is a major challenge in plant growth, severely affecting development and yield. Enhancing drought tolerance is thus a highly desired goal for agriculture. Here, we report that the loss-of-function of two drought-induced genes, *GASA3* and *AFP1*, significantly enhances drought tolerance in *Arabidopsis thaliana*. While constitutive expression of *GASA3* and *AFP1* increased drought sensitivity compared to wild type (WT) plants, a *gasa3afp1* double mutant exhibited superior drought tolerance compared to the single mutants. Enhanced drought tolerance of *gasa3*, *afp1* and *gasa3afp1* is likely due to reduced water loss caused by smaller stomatal apertures and thus lower transpiration rates. Moreover, *gasa3* and *afp1* mutants accumulated higher levels of abscisic acid (ABA) under drought conditions than WT plants, concomitant with a stronger up-regulation of ABA-responsive genes such as *RD29A/B*, *ABF2/3*, and *ABI5*. The stronger ABA increase in the mutants seems to result from hydrolysis of abscisic acid-glucosyl ester (ABA-GE) from vacuolar stores via the β-glucosidase *BG2* rather than by *de-novo* biosynthesis. Promoter analysis revealed the presence of ABA-responsive and drought stress-related cis-acting elements within the *GASA3* and *AFP1* promoter regions. RT-qPCR confirmed that the expression of both genes increased under drought. However, *GASA3* induction was significantly reduced in the absence of *AFP1,* suggesting that *AFP1* is involved in the modulation of *GASA3* expression. Our findings identify a novel *AFP1/GASA3*-driven control circuit that negatively regulates drought tolerance by suppressing stomatal closure and attenuating ABA signalling.

## 1. Introduction

Plants as sessile organisms are exposed to constantly changing environmental conditions. Successful plant development and adaptation is thus determined by long-time genetically inherited programs that are fine-tuned by short-term responses to abiotic and biotic stresses. For that purpose, plants contain a number of intercommunicating signalling networks to coordinate their responses to various external and internal stimuli (Signorelli, 2022). These signalling pathways balance the often-contrary needs of growth, development and stress protection and thus modify the stress response accordingly (Claeys & Inzé, 2013; Verma *et al*., 2016). However, stress response often comes at the cost of reduced yield (Zhu, 2016). To select for traits that provide yield stability under environmental challenges, it is crucial to gain an in-depth understand of the mechanisms of stress response.

Drought stress caused by limited water availability is considered as one of the major abiotic stresses that negatively affects plant growth, development and reproductivity (Claeys & Inzé, 2013; Tenorio Berrío *et al*., 2022). Accordingly, plants have developed various physiological and morphological adaptations to reduce water loss and optimize water use efficiency. Resistance mechanisms to drought comprise a wide range of cellular processes including global reprogramming of transcription, post-transcriptional modification of RNA and post-translational modification of proteins, ultimately leading to adaptive alteration of metabolism and plant development (Yang *et al*., 2010).

ABA is an essential phytohormone that regulates plant adaptation to drought (Muhammad Aslam *et al*., 2022). Plant exposure to drought stress induces the elevation of the ABA content, leading to stomatal closure and activation of drought related genes (Nakashima *et al*., 2014). The biosynthesis of ABA starts in the plastids, from ß-carotene, leading to xanthoxin, which is transported to the cytosol. Xanthoxin is then converted to ABA-aldehyde, which is subsequently oxidized to ABA (Wu *et al*., 2023). The core ABA-signaling network consist of three major components: the ABA receptors PYRABACTIN RESISTANT/PYR1-LIKE/REGULATORY COMPONENT OF ABA RECEPTOR (PYR/PYL/RCAR), negative regulators in form of protein phosphatases 2C (PP2Cs), positive regulators such as sucrose non-fermenting 1-related protein kinases (SnRKs) and basic leucine zipper (bZIP) transcriptional activators such as ABA-INSENSITIVE 5 (ABI5) and ABI5 homologous ABA-RESPONSIVE ELEMENT BINDING FACTORS (ABFs), also known as AREBs for ABA-RESPONSIVE ELEMENT BINDING PROTEINS (Ali *et al*., 2020). In the absence of ABA, high PP2C activity results in deactivation of SNRK2 by dephosphorylation (Hirayama & and Umezawa, 2010). In the presence of ABA, a complex between PYR/PYL/RCAR and PP2CA is formed, and SNRK2 is activated by phosphorylation. Phosphorylated SNRK2 subsequently phosphorylates ABI5/ABFs which then bind to *cis-*elements in target gene promoters known as ABREs (ABA-responsive elements) to activate gene expression (Choi *et al*., 2000; Ali *et al*., 2020).

ABI5 BINDING PROTEINs (AFPs) interact with AREBs such as ABI5 and promote their degradation by E3 ubiquitin ligase, hence negatively regulating ABA signalling (Lopez-Molina *et al*., 2003; Wei *et al*., 2022; Vittozzi *et al*., 2024). In Arabidopsis, four AFPs have been identified and AFP1 and its close homolog AFP2 have been shown to repress bZIP activation of certain ABRE-regulated genes (Lynch *et al*., 2022). Moreover, the *MEDIATOR-OF-OsbZIP46-DEGRADATION-AND-DEACTIVATION* (MODD), an ortholog of AFP3 in rice, was shown to be involved in negative regulation of drought tolerance (Tang *et al*., 2016). The *GIBERELLIC ACID STIMLATED ARABIDOPSIS* (*GASA*) family in Arabidopsis comprises genes with homology to *GA-STIMULATED TRANSCRIPT 1* (*GAST1*) from tomato (*Solanum lycopersicum*). *GASA* genes encode low-molecular-weight peptides, also called SNAKINs, that have been shown to play various roles in plant development as well as plant stress regulation (Bouteraa *et al*., 2023). Of the 14 *GASA* family members in Arabidopsis, *AtGASA4* has been shown to positively regulate heat stress tolerance (Ko *et al*., 2007), whereas *AtGASA5* has been demonstrated to negatively regulate thermotolerance (Zhang & Wang, 2011). Furthermore, *AtGASA14* has been shown to positively regulate salt stress tolerance by reducing ROS accumulation (Sun *et al*., 2013). *AtGASA3* so far has only been proposed to have increased transcript levels in seeds during dessication (Aubert *et al*., 1998), without further characterization of any role during abiotic stress, particularly drought.

In this study, we showed that the loss-of-function of two drought-induced genes, *GASA3* and *AFP1,* led to a strong increase in drought tolerance of *Arabidopsis thaliana.* Phenotypic analyses under drought conditions showed that while *s*ingle as well as *gasa3afp1* double mutants have enhanced drought tolerance, the constitutive overexpression lines have a reduced tolerance compared to WT plants. Expression of *GASA3* and *AFP1* is induced by drought and ABA according to RT-qPCR, however, induction of *GASA3* remains rather low in the absence of *AFP1*. Furthermore, we detected a reduced water loss most likely caused by smaller stomatal apertures and thus transpiration rates in *gasa3* and *afp1* single mutants as well as in *gasa3afp1*double mutants, suggesting an involvement of these two genes in supressing stomatal closure. Additionally, we show that *gasa3* and *afp1* plants accumulate higher levels of ABA under drought conditions than WT, concomitant with a further increase in the expression of ABA-responsive genes. Increased expression of the vacuolar β-glucosidase *BG2* and repression of genes involved in ABA synthesis suggest that the elevated ABA levels are caused by hydrolysis of abscisic acid-glucosyl ester (ABA-GE). Overall, our results indicated that *GASA3* and *AFP1* are negative regulators of drought stress tolerance in Arabidopsis via the ABA-signalling pathway, with *AFP1* involved in the modulation of *GASA3* expression.

## 2. Materials and Methods

### 2.1. Plant material and growth conditions

This study was performed using the Columbia ecotype of *Arabidopsis thaliana* (Col-0) and all transgenic lines were generated in this background. *gasa3* (SAIL_198_A11) and *afp1* (SAIL_13_C02) T-DNA insertion mutants were obtained from Nottingham Arabidopsis Stock Center (NASC, UK). The *aba2-1* mutant (Cheng *et al*., 2002; Lin *et al*., 2007) was a kind gift from Prof. Wan-Hsing Cheng, University of Taiwan. For most experiments, plants were directly placed into standard plant potting soil pre-treated with Confidor WG 70 (Bayer Agrar, Germany). For some experiments, sterilized seeds were sown on ½ MS (Murashige and Skoog medium, Duchefa Biochemie, The Netherlands) plates with 1% (w/v) sucrose and 0.6% (w/v) phytagel. Plants were grown in climate-controlled rooms under long day conditions (LD; 16h light / 8 h dark) with a light intensity of 100 µmol photon*m^-2^*s-^1^ (Philips TLD 18W lamps of alternating 830/840 light temperature).

### 2.2. Generation of transgenic lines

For the generation of 35S::*GASA3*-*YFP* or 35S::*AFP1*-*YFP* lines, the coding sequences of *GASA3* and *AFP1* without the stop codon were cloned into the pBIN19 vector (Datia *et al*., 1992) in frame with the coding region of *YFP*. The resulting expression cassettes were stably inserted into the genome of Col-0 plants using *Agrobacterium tumefaciens* and floral dipping (Zhang *et al*., 2006). Two independent T3 lines were selected by BASTA resistance, and constitutive expression was confirmed by RT-qPCR. Two *gasa3afp1* double mutant lines originating from independent crosses were generated by crossing homozygous single mutant lines. All primers used for cloning and screening are listed in Supplementary Table S1.

### 2.3. Plant phenotyping

For phenotyping of the different plant lines under drought stress, seeds were first germinated in batches on soil for a week. Subsequently, the seedlings were transferred into single pots filled with 100 grams of soil and kept well-watered until day 18. Before discontinuation of watering, individual pot weights were measured and all other pots were set to the pot with the highest weight using tap water. Plants were exposed to progressive drought or well-watered control conditions (50 ml of tap water each other day) for up to 12 days. The position of the plant pots was randomized throughout the experiments to avoid positional effects on the plant growth.

Two crucial parameters were closely monitored: Soil Water Content (SWC) and Real Leaf Water Content (RWC) of the rosette leaves. For measuring the soil water content as a percentage, the following formula was used: {(pot weight during measurement) − (empty pot weight)}/ {(initial pot weight) − (empty pot weight)} × 100. RWC was determined using previously published protocols (Barrs & Weatherley, 1962; Bouchabke *et al*., 2008). by measuring three different weights from whole rosettes (without reproductive tissue): the fresh weight (FW), the turgid weight (TW, after submerging the rosette in water overnight), and the dry weight (DW; measured after drying the rosettes at 72°C for 3 days). The formula applied for RWC expressed as percentage was: (FW-DW)/(TW-DW)x100.

### 2.4. Gene induction analysis

The expression of *GASA3* and *AFP1* was investigated after treatment with different compounds. For that purpose, 8 ml of either 100 µM ABA (Sigma-Aldrich, USA),100 µM methyl jasmonate (MeJA, SERVA, Germany), 100 µM gibberellic acid (GA3, Duchefa Biochemie, Netherlands) or 20 % (w/v) polyethylene glycol (PEG)-6000 (Carl Roth GmbH, Germany) were applied directly onto ½ MS plates with 21-day old plants and incubated for 0, 1, 3, 6, 9 or 24 hours under LD conditions. Whole seedlings were frozen using liquid nitrogen, ground into a fine powder and used for total RNA extractions as describe below in 2.7.

### 2.5. Stomatal measurements and estimation of transpiration rate

Stomatal aperture was quantified following a modified version of a previously established protocol (Eisele *et al*., 2016). Briefly, 7^th^ or 8^th^ leaves of 32-day old plants grow on soil were incubated for 2 hours with imaging buffer (10 mM MES, pH 6.15, 5 mM KCl, 50 µM CaCl2). Epidermal peels were carefully separated from the mesophyll and fixed to a glass slide using medical adhesive tape. Images were taken under Bright Field settings using a Leica SP8 Lightning using the integral LAS X software and were further processed using the Fiji/ImageJ software (Schindelin *et al*., 2012). For estimation of stomatal density, the diameter of the field of view (FV) was calculated (πr^2^) and used to normalize the count. Transpiration rate (mmol*m^-2^*s^-1^) was quantified using a LiCOR LI 6000 porometer/fluorometer (LI-COR Environmental GmbH, Germany).

### 2.6. Quantification of ABA

For ABA measurements, whole plant rosettes were frozen in liquid nitrogen, and ground into a fine powder. The extraction and quantification followed a previously established protocol (Pan *et al*., 2010). Briefly, 50 mg of rosette tissue were harvested and immediately frozen in liquid nitrogen and homogenized with a pre-cooled mortar and pestle. Samples were handled using only liquid nitrogen throughout harvesting ensuring minimal damage due to repetitive freeze-thawing. This was followed by the addition of 500 µl extraction solvent (2-propanol/water/conc. HCl in a ratio 2:1:0.002, v/v/v) and 25 ng D6-ABA, addition of 1 ml dichloromethane and phase separation, removal of the lower phase, and nitrogen assisted drying of the upper phase. The dried matter was resuspended in 0.1 ml of methanol:0.1% formic acid in water (1:1, v/v). Phytohormones were separated on a reverse phase C18 Gemini HPLC column and analysed using a QTRAP 6500+ LC-MS/MS system (Sciex, Germany). Data evaluation was carried out using the MultiQuant^TM^ 3.0.2 software (Sciex, Germany). The concentrations of ABA were determined relative to the internal standards, and expressed as ng/g F.W. All used solvents were of HPLC grade or LC-MS grade.

### 2.7. Estimation of total anthocyanin content

Anthocyanins were quantified according to a previous protocol (Nakata & Ohme-Takagi, 2014). The rosettes of 32-day old plants (control and drought) were flash frozen using liquid nitrogen and pulverized into a fine powder. Based on the fresh weight of the samples, approximately 5 volumes of extraction buffer (45 % methanol, 5 % acetic acid) were added and vortexed thoroughly. The mixture was centrifuged two times at 12000g for 5 min and absorbances of the supernatants were recorded at 530 and 637 nm. The amount of anthocyanin per gram fresh weight (g^-1^*F.W.^-1^) was calculated by the formula: (Abs530/g F.W.) = [Abs530 - (0.25 x Abs637)] x 5.

### 2.8. RNA-extraction, cDNA synthesis and RT-qPCR

For RNA extraction of soil-grown plants, whole rosettes were harvested and ground into a fine powder using liquid nitrogen. RNA was extracted from 100 mg of this powder using the Roboklon Plant RNA Kit (Roboklon GmbH, Berlin, Germany). The quality of the RNA was assessed using a Nanodrop Spectrophotometer or by separation on a 1% agarose gel. cDNA synthesis was carried out from at least 500 ng RNA using the Revert Aid First strand cDNA kit (ThermoFisher Scientific, USA) and oligodT18 primers. The reaction was carried out for 1 hour at 42 °C, followed by termination by heating at 72 °C for 10 min.

RT-qPCR was carried out on a Bio-Rad CFX96 touch system (Bio-Rad Laboratories, Germany). Gene expression data were analyzed using the 2^–ΔΔCt^ method (Livak & Schmittgen, 2001) and normalized to the geometric means of two reference genes: *AtACT2* and *AtTUB2* (Vandesompele *et al*., 2002). All primers used are listed in Supplementary Table S1. Unless otherwise mentioned in the figure legends, for RT-qPCR analyses, 32-day-old plants either under control or drought conditions were used.

### 2.9. Statistical analyses

Statistical analyses were conducted using R version 4.3.2 (R Core Team, 2023; https://www.r-project.org/). A two-tailed Student’s *t*-test (*P* < 0.05) was used to compare drought and control samples, employing the base t.test() function. For datasets involving multiple groups or treatments, one-way or two-way ANOVA was performed, followed by Tukey’s HSD test (*P* < 0.05), using the R packages agricolae, tidyverse, and ggplot2. For two-way ANOVA both capital and small letters were used, where the capital letters signified variance due to treatment (ABA or drought) and the small letters depicted variance due to genotypic differences.

## 3. Results

### 3.1. *GASA3* and *AFP1* expression is strongly induced by progressive drought

While screening for drought-responsive genes using the RNA-seq dataset of a recent study in *Arabidopsis thaliana* (Mahmud *et al*., 2022), we identified two highly drought induced genes: *GASA3* (AT4G09600) and *AFP1* (AT1G69260). *GASA3* belongs to the gibberellic acid-stimulated (GAST) Arabidopsis family implicated in a wide range of functions like plant growth, development and fruit ripening (Vittozzi *et al*., 2024; Bouteraa et al., 2023). *AFP1*, on the other hand, is best studied during germination and has been shown to promote the degradation of the bZIP transcription factor ABI5, which is known to promote the expression of ABA-responsive genes (Lopez-Molina *et al*., 2003; Wei *et al*., 2022).

We thus analysed the expression of *GASA3* and *AFP1* during a time-course of 14 days of progressive drought compared to well-watered plants. As before (Mahmud et al., 2022), water withholding was started when soil-grown plants were 18 days old. RT-qPCR showed that under well-watered conditions the expression of *GASA3* and *AFP1* was very low and showed not significantly changes during the course of the experiment (Figure 1a). By contrast, expression of *GASA3* and *AFP1* increased with a fold change (FC) of about 10 after 6 and 7 days of water withholding, respectively. Expression of *AFP1* further increased gradually to a FC>60, while *GASA3* expression showed an exponential increase to an FC>1000 on day 12 and >2000 on day 14 (Figure 1a). These results confirm the RNA-Seq data from the previous study (Mahmud *et al*., 2022) but also show that onset of gene induction occurs early on after water withholding.

**Figure 1:**
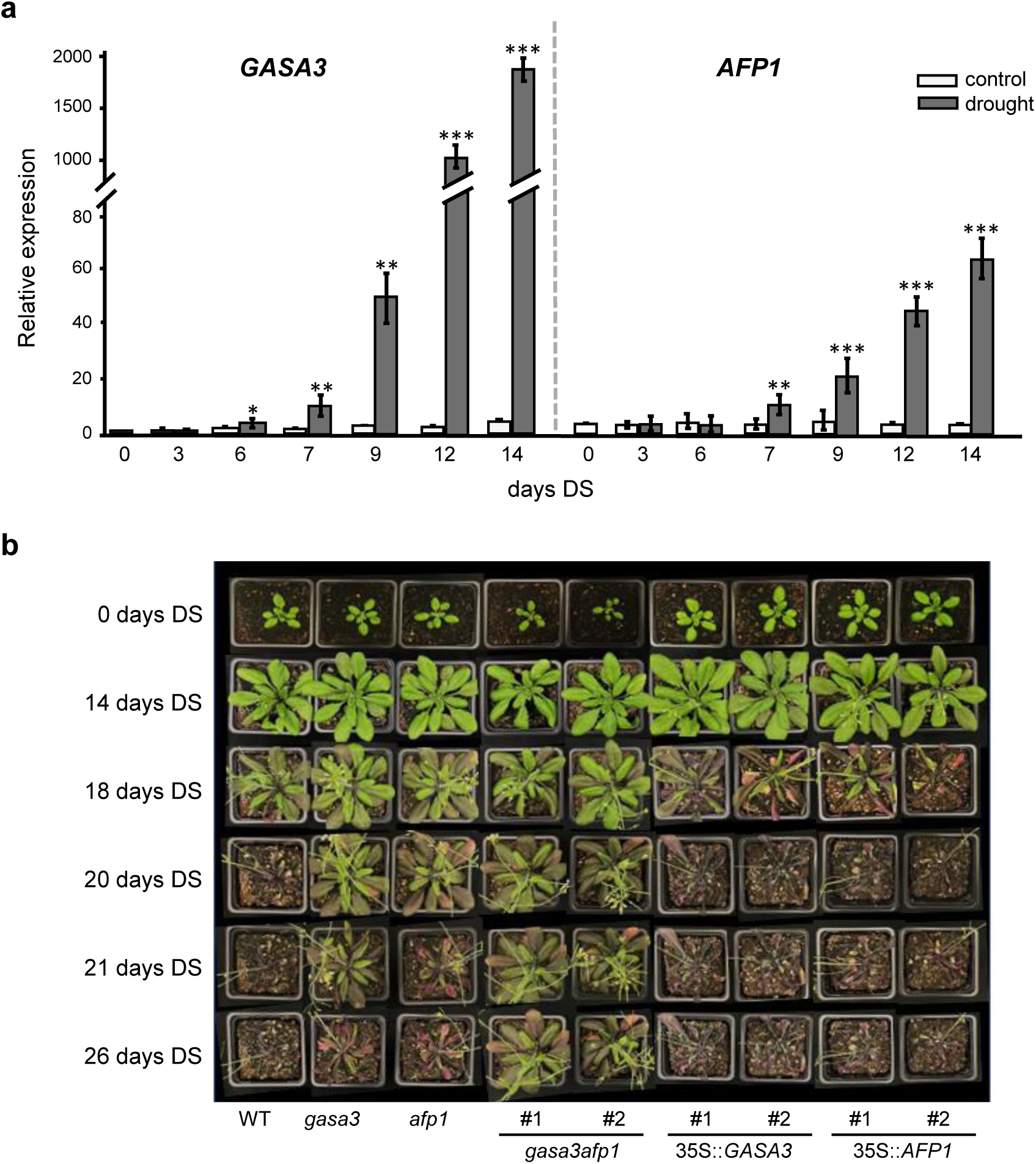
Effect of *GASA3* and *AFP1* on drought tolerance in *Arabidopsis thaliana*. (a) Relative expression of *GASA3* and *AFP1* in WT plants at various days of progressive drought stress (DS) on soil. Water withholding was started when the plants reached an age of 18 days. Data represent means ± SE of three independent biological repeats (n=3). Statistical analyses were carried out using two-tailed student T-test between drought and control (*P<0.05, **P<0.01, ***P<0.001). (b) Drought stress phenotype of WT, *gasa3* and *afp1* single mutants, a *gasa3afp1* double mutant, and lines expressing 35S::*GASA3* and 35S::*AFP1* in WT background at different days of progressive drought stress (DS). The images are representative of several individual experiments

### 3.2. *GASA3* and *AFP1* negatively regulate drought tolerance in Arabidopsis

To evaluate the effect of *GASA3* and *AFP1* on drought tolerance, we analysed the growth phenotype of homozygous T-DNA insertion lines for *gasa3* and *afp1*, a *gasa3afp1* double mutant and lines expressing YFP-tagged *AFP1* and *GASA3* under control of the 35S promoter in the WT background, which we refer to as 35S::*GASA3* and 35S::*AFP1* (Supplementary Figure 1). RT-qPCR analyses confirmed complete lack of expression of the respective gene in *gasa3* and *afp1* (Supplementary Figure 1b) and constitutively elevated expression in 35S::*GASA3* and 35S::*AFP1* (Supplementary Figure 1d). No difference in growth was observed compared to WT up to 14 days of water withholding (Figure 1b). However, clear differences could be observed upon longer drought periods. While the WT showed signs of wilting on day 18 and was nearly completely wilted on day 20, *afp1* plants only showed strong wilting on day 21 and *gasa3* plants on day 24. Plants from the 35S::*GASA3* and 35S::*AFP1* lines wilted a bit earlier than the WT plants, while the double mutant lasted even longer than the single mutants. All in all, these results show that despite being induced under drought, both *GASA3* and *AFP1* have a negative impact on drought tolerance, which to a certain degree is additive.

### 3.3. *GASA3* and *AFP1* affect water loss through modulation of the stomatal aperture

Water loss through transpiration is an important factor related to drought tolerance. To, determine the rate of water loss, the RWC of the rosette leaves was measured in the different lines on day 14 of drought, when all plants still looked similarly healthy, and on day 18, when differences in the drought tolerance was clearly visible (Figure 1b). Already on day 14 the lines showed a difference in RWC, with the single and double mutants still retaining more than 80 % RWC, while RWC in the 35S::*GASA3* and 35S::*AFP1* lines dropped to around 60% (Figure 2a). On day 18, single and double mutants still retained RWCs of over 80 %, while RWC dropped below 60% in the wild type and below 30% in the 35S::*GASA3* and 35S::*AFP1* lines (Figure 2a).

**Figure 2:**
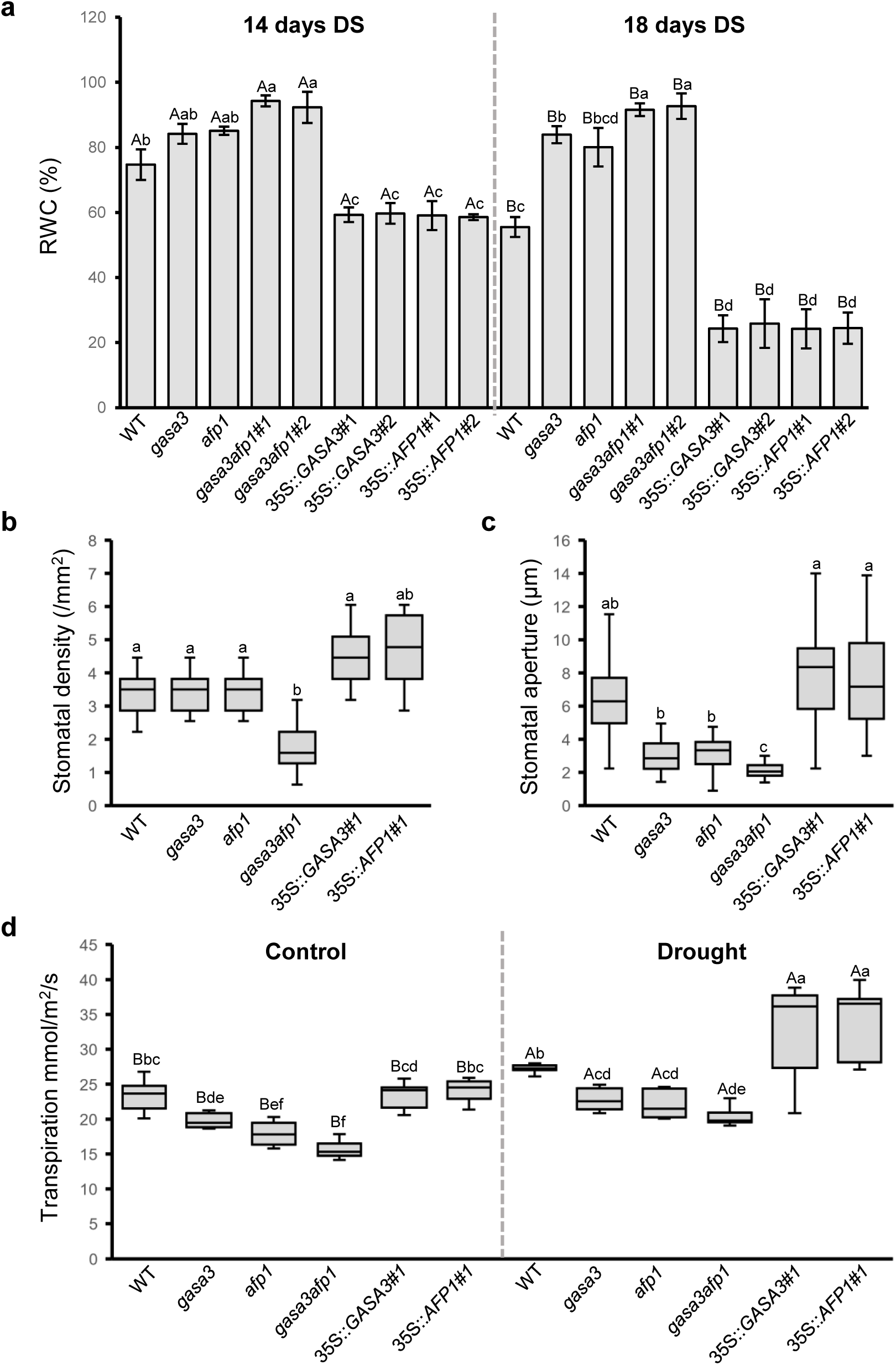
Effect of *GASA3* and *AFP1* on stomata regulation and leaf relative water content. (a) Leaf relative water content (% RWC) of plants at days 14 and 18 of progressive drought stress (DS). Data represents means ± SE of three independent replicates (n=3). Assessment of stomatal density and (c) stomatal aperture measured on leaves No. 7 and 8 of plants grown under control conditions for 32 days. For stomatal density, each replicate quantified leaves from two individual plants. For stomatal aperture, each replicate quantified 20 stomata in leaves from two individual plants. (d) Transpiration rates in leaves of 32 day-old plants grown under control and drought conditions. Data represent means ± SE from three biological replicates (n = 3). For all measurements the statistics were carried out using ANOVA and Tukey‘s Post-Hoc HSD tests (P<0.05).

The number of stomata per leaf area as well as their apertures determines the transpiration rate. While the stomata number is a fixed trait determined during development, stomata aperture is regulated dynamically in response to various parameters and the process encompasses multiple signaling pathways (Araújo *et al*., 2011). With regards to stomata density, a discernible difference was only observed in the double mutant plants, which possess significant fewer stomata per mm (Figure 2b). By contrast, stomatal aperture in the different plants was very much in line with the observed phenotype, with the smallest aperture observed in the single and double mutant and the largest in the 35S::*GASA3* and 35S::*AFP1* lines (Figure 2c). This is also in line with the observation that loss of *GASA3* or *AFP1* increases the transcript levels of two genes associated with stomatal closure, the beta-thioglucoside glucohydrolase *TGG1* (Islam *et al*., 2009) and the slow (S)-type anion channel *SLAC1* (Deng *et al*., 2021), even under control conditions (Supplementary Figure S2a). Furthermore, in the leaves of *gasa3*, *afp1* and *gasa3afp1* the transpiration rates were lower than those in the WT and the 35S::*GASA3* and 35S::*AFP1* lines, under drought as well as under control growth conditions (Figure 2d). Together, these results suggest that regulation of stomata aperture is the cause behind the alteration in transpiration and thus the better drought tolerance of the *gasa3* and *afp1* single and *gasa3afp1* double mutant plants.

### 3.4. Expression of *GASA3* and *AFP1* depends on ABA

In silico analysis of the *GASA3* and *AFP1* promoter regions (-1kb) shows the presence of various cis-elements known to confer response to plant hormones and abiotic stresses (Supplementary Figure S3). These include the Abscisic Acid Responsive Element (ABRE), of which a single one was detected in the *GASA3* promoter region and four in the *AFP1* promoter region.

In order to examine whether ABA regulates the expression of *GASA3* and *AFP1*, we analysed WT seedlings grown for 21 days on ½ MS phytagel plates that were treated with exogenous ABA. We also included MeJA and GA3 since cross-talk between ABA and jasmonate has been described in drought response (de Ollas & Dodd, 2016; Mahmud et al. 2022) and GASA proteins were originally identified in relation to GA signaling (Shi *et al*., 1992). We first treated the seedlings with 20% PEG to confirm that both genes are also upregulated under these growth conditions when drought is mimicked (Figure 3a and b). Moreover, addition of 100 µM ABA increased the expression of *GASA3* and *AFP1*, while neither MeJA nor GA3 had any inducing effect. A similar picture emerged when gene expression was analysed separately in roots and shoots indicating that drought and ABA induction of *GASA3* and *AFP1* is not specific for photosynthetic tissues (Supplementary Figure S4).

**Figure 3:**
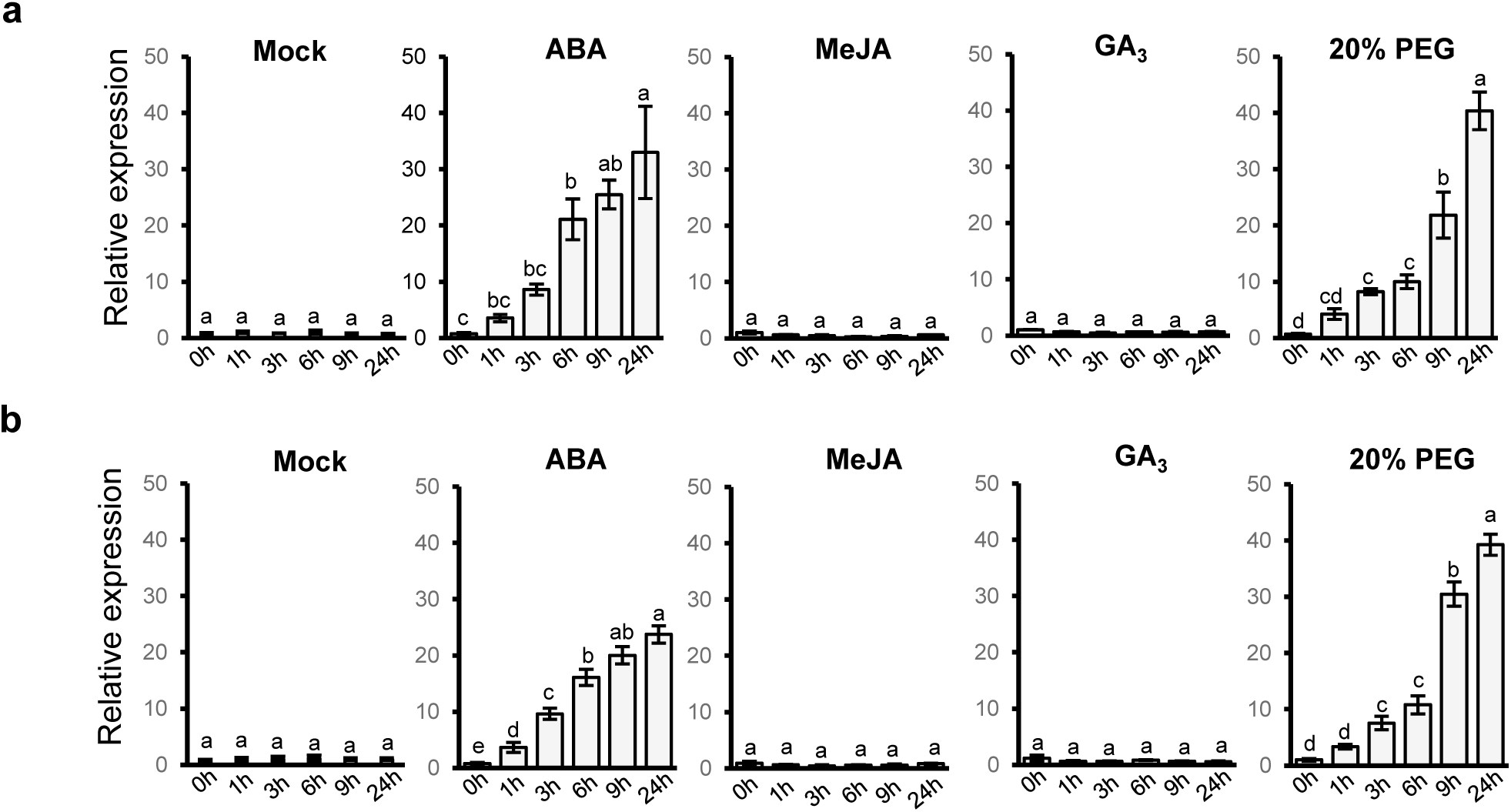
Induction of *GASA3* and *AFP1* expression by PEG and various hormones. Relative expression of (a) *GASA3* and (b) *AFP1* in 21-day old WT seedlings grown on ½ MS plates treated with either ddH_2_O, 100 µM ABA, 100 µM MeJA, 100 µM GA_3_ or 20% PEG-(a) 6000. Data represent means ± SE of three independent biological replicates (n=3). Statistical analyses were performed with one-way ANOVA and Tukey‘s Post-Hoc HSD tests (P<0.05).

We furthermore analysed the transcript levels of *GASA3* and *AFP1* under progressive drought in the *aba2-1* mutant, which is impaired in ABA biosynthesis (Cheng *et al*., 2002). Compared to WT, no induction of either *GASA3* and *AFP1* could be observed in *aba2* (Figure 4a and b) but induction was restored by addition of external ABA (Figure 4c and d). These results support a direct role of ABA in the drought-induction of *GASA3* and *AFP1*.

**Figure 4:**
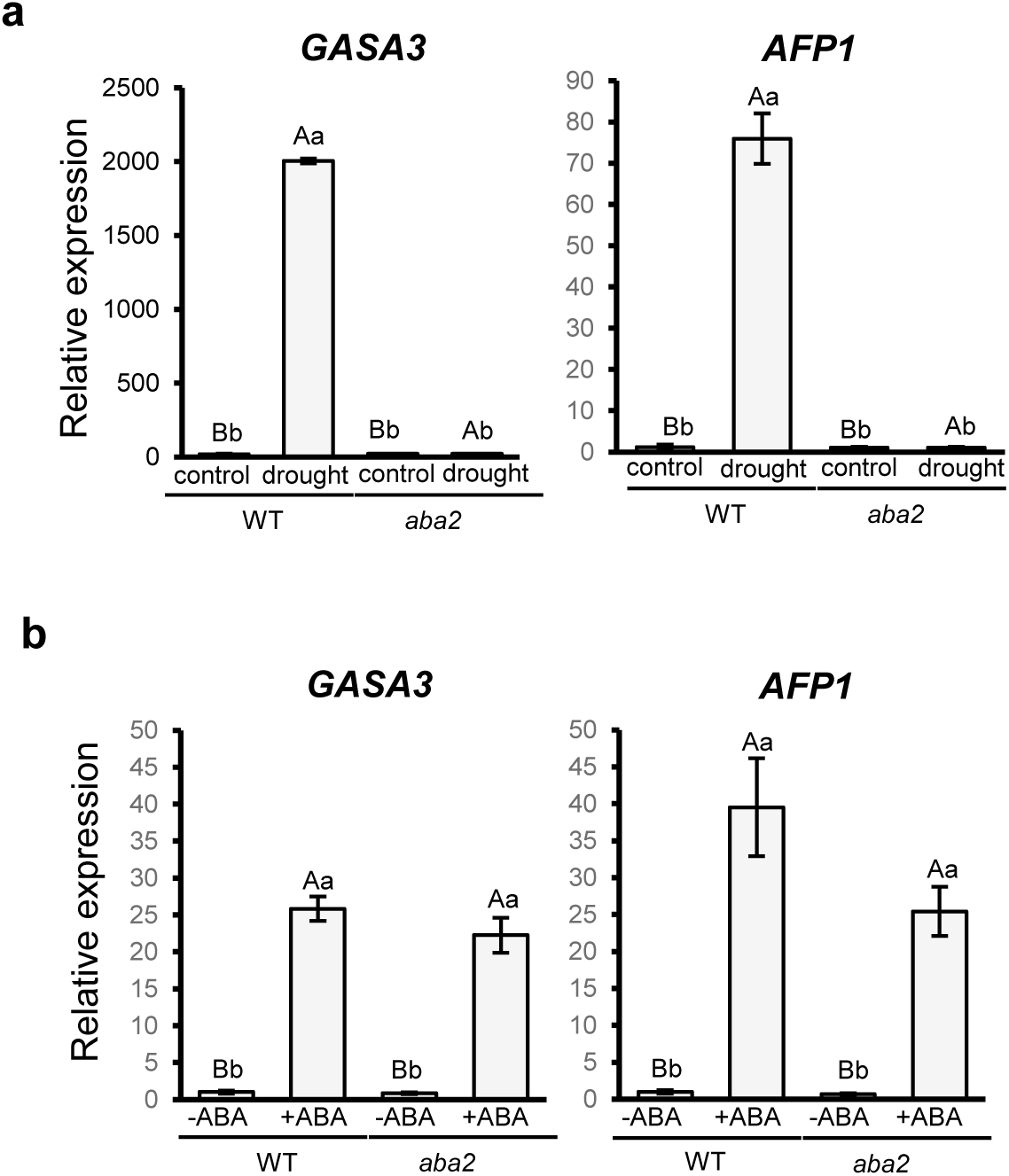
ABA-dependency of *GASA3* and *AFP1* expression. (a) Relative expression of *GASA3* and *AFP1* in WT and *aba2* mutant plants grown on soil under control and progressive drought conditions. (b) Relative expression of *GASA3* and *AFP1* in 21-day old WT and *aba2* seedlings grown on ½ MS plates treated with either ddH_2_O or 100 µM ABA for 24 hours. Data represent means ± SE of three biological replicates and statistical analyses were carried out with two-way ANOVA along with Tukey‘s Post-Hoc HSD tests (P<0.05).

### 3.5. *GASA3* and *AFP1* affect ABA signaling via release of ABA-GE

As described (Mahmud *et al*., 2022), we found an elevated ABA content in the WT plants after 14 days of drought (Figure 5a). However, the increase in ABA content was stronger in the *gasa3* and *afp1* mutant lines. Analysis of the expression of several genes related to ABA in *gasa3, afp1* and the WT (Figure 5b-d) showed that the expression of ABA-responsive genes, such as *ABF2*, *ABF3*, *RD29A,* and *RD29B,* was much stronger induced in *gasa3* and *afp1* compared to WT (Figure 5d). By contrast, the drought induction of *PP2CA*, which forms an important negative feed-back loop of ABA response, is supressed in the mutants (Figure 5d). Similarly, the expression of ZEP/*ABA1* and *ABA2*, whose gene products catalyse key steps in ABA biosynthesis (Chen *et al*., 2020), was induced under drought in the WT but suppressed in *gasa3* and *afp1* (Figure 5b), suggesting that the mutants do not produce the surplus ABA by *de-novo* biosynthesis from ß-carotene under drought. ABA can also be generated by activation of ABA-GE stored in the endoplasmic reticulum and vacuole via the β-glucosidases BG1 and BG2 respectively (Xu *et al*., 2012; Han *et al*., 2020). We observed an up-regulation of *BG2* but not *BG1* in both mutants under drought, a response that is absent in WT plants (Figure 5c). These findings suggest that the increased ABA levels in *gasa3* and *afp1* derive from conjugated ABA-GE stored in the vacuole.

**Figure 5:**
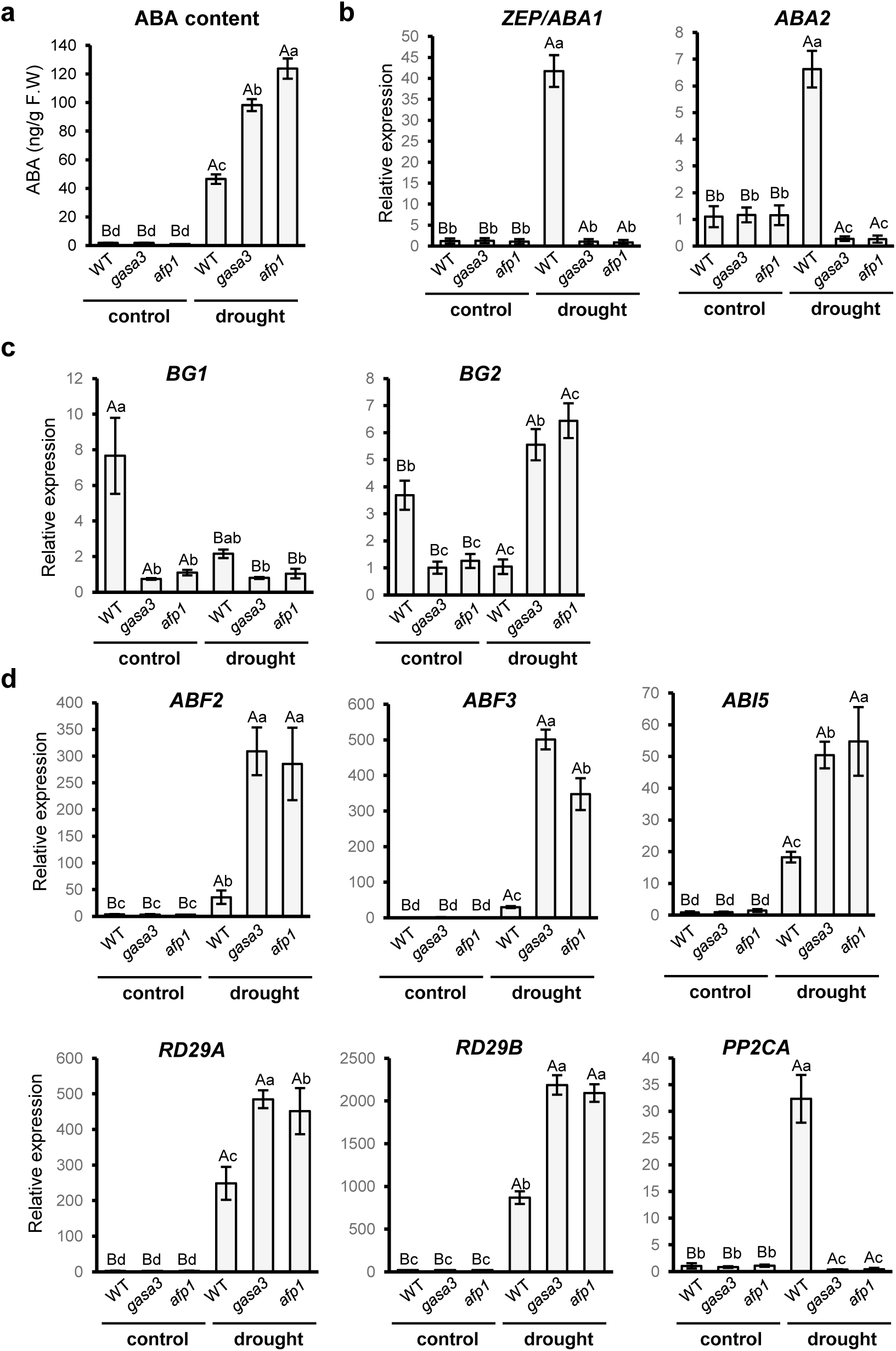
Effect of *GASA3* and *AFP1* on ABA biosynthesis and the ABA-mediated drought response. WT, *gasa3* and *afp1* mutant plants grown under control and progressive drought conditions were investigated for (a) Endogenous ABA content of rosette tissue as well as relative expression of genes involved in (b) ABA biosynthesis, (c) ABA-GE activation, and (d) ABA-responses. Data represent means ± SE of three independent replicates (n=3). Statistical significance was estimated with two-way ANOVA and Tukey‘s HSD analyses (P<0.05).

### 3.6. *AFP1* acts as upstream regulator of *GASA3*

Our data so far raise the question, whether *GASA3* and *AFP1* function in the same drought response pathway. To address this question, we investigated the expression *of AFP1* in the *gasa3* mutant and *vice versa* (Figure 6a and b). Upon progressive drought, *AFP1* was induced in the *gasa3* mutant line to a level even a bit higher than in WT (Figure 6a) but *GASA3* induction was strongly reduced in the *afp1* mutant (Figure 6b). To confirm that the reduce GASA expression is indeed caused by a lack of *AFP1*, we introduced the 35S::*AFP1* construct into the *afp1* mutant background. This resulted in a low constitutive expression of *AFP1* under control conditions and a similar drought sensibility as the WT (Figure 6c). At the same time, strong induction of *GASA3* under drought was restored (Figure 6d). These data suggested that *AFP1* positively modulates *GASA3* expression under drought stress and that *GASA3* might be the key effector that drives drought susceptibility.

**Figure 6:**
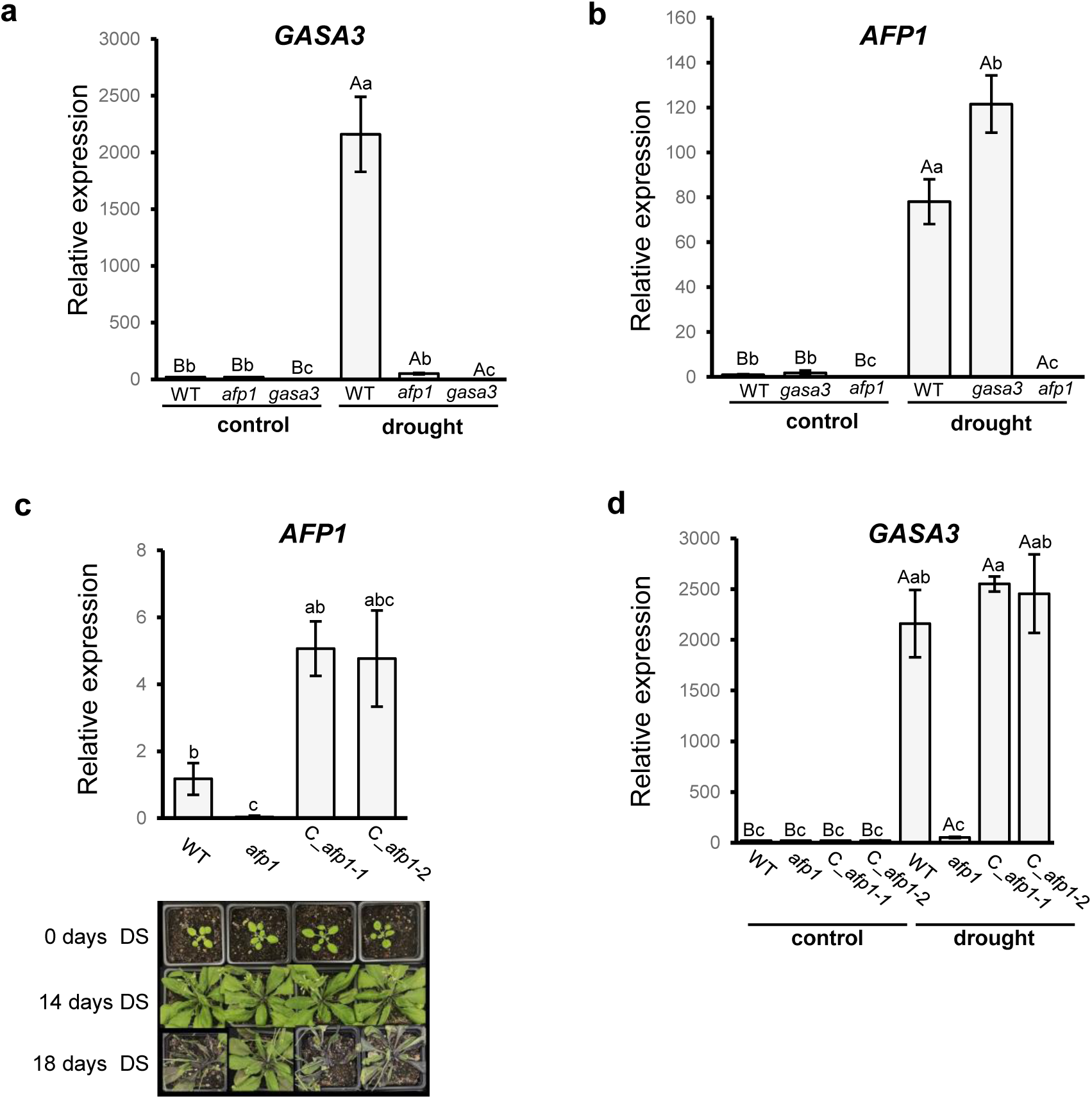
Role of *AFP1* in the expression of *GASA3*. Relative expression of (a) *GASA3* and (b) *AFP1* in WT, *gasa3* and *afp1* mutant plants grown under control and progressive drought conditions. (c) Relative expression of *AFP1* and drought phenotype of WT, *afp1* and two lines expressing 35S::*AFP1-YFP* in the *afp1* mutant background (C_*afp1*). DS: drought stress. The images are representative for several individual experiments. (d) Relative expression of *GASA3* in WT, *afp1* and C_*afp1* lines grown under control and progressive drought conditions. RT-qPCR data represent means ± SE of three biological replicates (n=3), where statistical analyses were carried out using two-way ANOVA (time period of drought and genotype) and Tukey‘s Post-Hoc HSD tests (P<0.05).

## 4. Discussion

Plants have evolved various cellular and molecular mechanisms that enhance their acclimation to drought stress. In this study, we investigated the roles of *GASA3* and *AFP1* in the drought stress response, revealing a partial interdependent relationship between these two genes, in which *GASA3* is a downstream component of *AFP1* mediated signalling.

*GASA3* and *AFP1* exhibit a drought-dependent increase in transcript levels (Figure 1a) and loss-of-function mutants showed that *GASA3* and *AFP1* are negative regulators of drought tolerance in Arabidopsis (Figure 1b). Moreover, the *gasa3afp1* double mutants displayed a further enhanced drought tolerance, suggesting an at least partial additive function of GASA3 and AFP1. Our data further suggests that *GASA3* and *AFP1* negatively regulate drought tolerance through a mechanism that is primarily driven by stomatal movement rather than differences in stomata development. However, differences in other drought related traits not analysed in this study might affect the drought phenotype of the different lines. These could include traits pertaining to leaves as well as roots, since *AFP1* and *GASA3* are induced by drought in both tissues.

While *GASA3* and *AFP1* expression was induced by ABA (Figure 3), the ABA content was significantly increased under drought stress in *gasa3* and *afp1* plants compared to WT. This suggests that *GASA3* and *AFP1* might be part of a negative feedback loop, regulating ABA biosynthesis in Arabidopsis (Figure 7). However, *de-novo* biosynthesis of ABA seems to be rather supressed in the absence of *GASA3* and *AFP1*. Instead, increased expression of *BG2* suggests that the mutants generate ABA from conjugated ABA-GE stored in the vacuole. Consistent with the higher accumulation of ABA, *gasa3* and *afp1* plants showed an up-regulation of core ABA-responsive genes such as the ABREs *ABF2, ABF3, and ABI5*, which are crucial regulators of the ABA-induced transcriptional network (Choi *et al*., 2000; Vittozzi *et al*., 2024), or *RD29A*, a key component of ABA mediated drought responss (Msanne *et al*., 2011; Jia *et al*., 2012). g*asa3* and *afp1* plants also show an increase in anthocyanin production during drought that may contribute to their enhanced drought tolerance (Supplementary Figure 3c) since anthocyanins function as ROS-scavenging antioxidant and several studies have revealed a positive link between anthocyanin levels and drought tolerance in Arabidopsis (Nakabayashi *et al*., 2014). Overall, our finding fit well into current models on the role of ABA in drought response (Figure 7). The higher ABA level observed in the g*asa3* and *afp1* mutants would result in an increased phosphorylation of SnRK2, which then phosphorylates ABREs, resulting in increased expression of ABA-responsive genes. These include the S-type anion channel *SLAC1* that contributes to stomata closure, while expression of its counterplayer *KAT-1* is repressed (Takahashi et al., 2017). SnRK2 also phosphorylates both SLAC1 and KAT-1, leading to an activation of the former and inhibition of the latter, which ultimately results in stomata closure.

**Figure 7:**
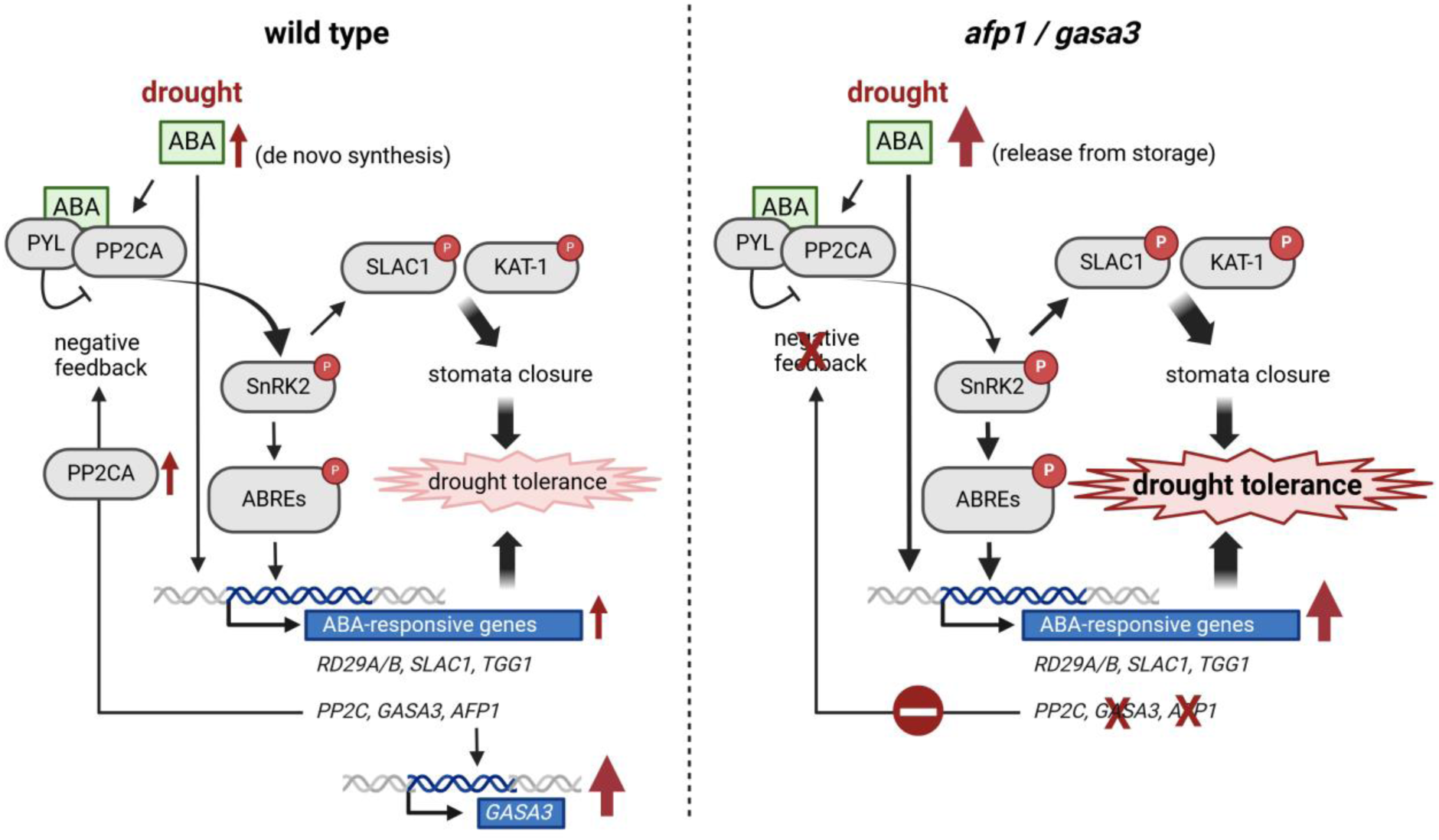
Model of ABA regulation of drought tolerance in WT compared to *gasa3* and *afp1* mutants. ABA-dependent protein phosphorylation and transcriptional regulation of ABA-responsive genes are at the core of ABA-dependent drought response. Increase in ABA synthesis and inhibition of the PP2CA negative feedback loop in the absence of *gasa3* and *afp1* ultimately result in increased expression of certain ABA-responsive genes as well as stomata closure via SLAC1 and KAT1 phosphorylation. Red arrows indicate changes in content of ABA or transcripts upon drought. Created in https://BioRender.com

Loss of *afp1* and *gasa3* moreover reduces the ABA-dependent induction of *PP2CA*, thereby preventing the negative feedback on SnRK2 (and thus SLAC1 and KAT-1) phosphorylation, further enhancing the effect of the increased ABA content (Figure 7).

In our study, we observed little difference in drought tolerance and related traits (RWC, stomata aperture etc.) between the *gasa3* and *afp1* mutants. The obvious reason is the lack of strong *GASA3* induction in the *afp1* mutant (Figure 6). While both genes can be induced by ABA, the expression of *GASA3* remains very low in the absence of *AFP1*. AFPs have been shown to bind to bZIP type transcription factors, thereby targeting them for proteasomal degradation (Lopez-Molina et al., 2003). *AFP1*-dependent increase in expression of *GASA3* could thus involve degradation of a *GASA3* repressor. Independent of the exact nature of this regulation, our data indicate that *GASA3* is the key effector and *AFP1* the regulator of the *AFP1*/*GASA3* dependent modulation of drought susceptibility (Figure 7). Further studies are needed to elucidate the exact mechanisms behind *GASA3* and *AFP1* dependent regulation in drought stress responses. This should include the expression of *GASA3* in the *afp1* mutant background driven by a drought induced promoter that is not regulated by *AFP1*. Since we could not observe a growth phenotype of the *afp1* and *gasa3* mutant under stress-free growth conditions and their expression under drought results in a reduced tolerance, their drought induction remains enigmatic. More studies are required that more closely resemble natural conditions including repeating cycles of mild drought and watering.

## Authors contribution statement

SB contributed to conceptualization, investigation (responsible for most experimental work), formal analysis (responsible for statistical analysis), validation, visualization, and writing - original draft as well as review & editing. BT, SL, DCR, and YS contributed to investigation (gene expression, phenotyping, promoter analysis). KG and PD contributed to investigation (hormone measurements) and writing - review and editing. FC contributed to conceptualization, formal analysis, validation, visualization, supervision, and writing - original draft as well as review & editing. UCV contributed to conceptualization, validation, visualization, funding acquisition, project administration, supervision, and writing - review & editing. All authors contributed to the article and approved the submitted version.

## Conflict of Interest

The authors have no conflicts to declare.

## Funding

This research was supported by DFG grant INST 217/939-1 FUGG to UCV.

## Data availability

The data that support the findings of this study are available from the corresponding author upon reasonable request.

## Supporting information

Table S1, Figure S1, Fugure S2, Figure S3

## Notes

### Competing Interest Statement

The authors have declared no competing interest.

