## Supplementary material for "Loss-of-function of the drought-induced genes *GASA3* and *AFP1* confers enhanced drought tolerance in *Arabidopsis thaliana*": Table S1, Figure S1, Fugure S2, Figure S3

**Supplementary table S1: List of primers used in this study.**
Primers used for genotyping, RT-qPCR and cloning in this study. Fw- forward primer, Rev- reverse primer, LP- left primer, RP- right primer.

| Name | Gene ID | Sequence | Purpose |
|---|---|---|---|
| *GASA3*_LP | AT4G09600 | TCCCAAATTTAAAAATCCACG | Genotyping |
| *GASA3*_RP | AT4G09600 | TGTGGACTCTACTTTGGGTGG | |
| *AFP1*_LP | AT1G69260 | AGAGGCATCGTTACAACAACG | |
| *AFP1*_RP | AT1G69260 | ACGCATTGTAGGACCTGAGAC | |
| LB_SAIL | | GCCTTTTCAGAAATGGATAAATAGCCTTGCTTCC | |
| *GASA3*_Fw | AT4G09600 | TGATTGCGGTGGGAGATGCAAAGG | RT-qPCR |
| *GASA3*_Rev | AT4G09600 | AGCAAGGACAAAGGTGGTGGTTCC | |
| *AFP1*_Fw | AT1G69260 | GTCTCCGAGTTGGAAACTAAAGCG | |
| *AFP1*_Rev | AT1G69260 | TGTTGTGGCTGTGTAGTTGATGG | |
| *ABF2*_Fw | AT1G45249 | AGTTACAACGAAAGCAGGCAAGG | |
| *ABF2*_Rev | AT1G45249 | CCTCCTTGCAGAAGATTCCTCATC | |
| *ABF3*_Fw | AT4G34000 | GTTCTCAACCTGCAACACAGTGC | |
| *ABF3*_Rev | AT4G34000 | TCCAGGAGATACTGCTGCAACC | |
| *RD29A*_Fw | AT5G52310 | TGGACAAAGCAATGAGCATGAGC | |
| *RD29A*_Rev | AT5G52310 | AGGTTTACCTGTTACGCCTGGTG | |
| *RD29B*_Fw | AT5G52300 | ACTGATCCCACGCATAAAGGTG | |
| *RD29B*_Rev | AT5G52300 | CTCGTCGGAAAGTCTTCTTCGC | |
| *SLAC1*_Fw | AT1G12480 | ATGGCCAATTCGACGGATGTTC | |
| *SLAC1*_Rev | AT1G12480 | ACCACGCCACTGAGAACTTAAATC | |
| *ZEP/ABA1*_Fw | AT5G67030 | TGGGCTTGGTCCTCTGTCTTTC | |
| *ZEP/ABA1*_Rev | AT5G67030 | CACCAACTCTTCCTGGATGTGG | |
| *ABA2*_Fw | AT1G52340 | TTGCTGCTGCAAACGCGAATC | |
| *ABA2*_Rev | AT1G52340 | AGCGTTCGCTACATCATCAACCG | |
| *NCED3*_Fw | AT3G14440 | TCGAAGCAGGGATGGTCAACAG | |
| *NCED3*_Rev | AT3G14440 | GCTCGGCTAAAGCCAAGTAAGC | |
| *AAO3*_Fw | AT2G27150 | TATGGAGTTGGAGTCAGCGAGGTG | |
| *AAO3*_Rev | AT2G27150 | GCCTTGAACAAATGCTCCTTCGG | |
| *BG1*_Fw | AT1G52400 | GAGTATGCACGACGCCATTTGC | |
| *BG1*_Rev | AT1G52400 | TGGTGACGGGTCAAGTTGTTCTG | |
| *BG2*_Fw | AT2G32860 | ACGTCAAAGCAGCTAGACGATCC | |
| *BG2*_Rev | AT2G32860 | TGTGAGAGGGCGAAGAAACCAG | |
| *PP2CA*_Fw | AT3G11410 | TCCTCTCTCCGTAGATCACAAGCC | |
| *PP2CA*_Rev | AT3G11410 | GGCAAGAACTCCAAGAACCCTAGC | |
| *ACT2*_Fw | AT3G18780 | TGCCAATCTACGAGGGTTTC | |
| *ACT2*_Rev | AT3G18780 | CTTACAATTTCCCGCTCTGC | |
| *TUB2*_Fw | AT5G62690 | TAACAACTGGGCCAAGGGACAC | |
| *TUB2*_Rev | AT5G62690 | ACAAACCTGGAACCCTTGGAGAC | |
| *GASA3*_Fw | AT4G09600 | cttATGGCAATCTTCCGAAGTAC | Cloning |
| *GASA3*_Rev | AT4G09600 | cttAGGGCACTTGAGACGG | |
| *AFP1*_Fw | AT1G69260 | cttATGGCGGAAGCAAACGAG | |
| *AFP1*_Rev | AT1G69260 | cttTAAGAGATTAGAAGGAGAAGAAGT | |

| | | |
|---|---|---|
| *GASA3*ApaI | AT4G09600 | cttGGGCCCATGGCAATCTTCCGAAGTAC |
| *GASA3*NotI | AT4G09600 | cttGCGGCCGCCAGGGCACTTG |
| *AFP1*ApaI | AT1G69260 | cttGGGCCCATGGCGGAAGCAAACGAG |
| *AFP1*NotI | AT1G69260 | cttGCGGCCGCCTAAGAGATTAGAAGGAGAAGAAGT |

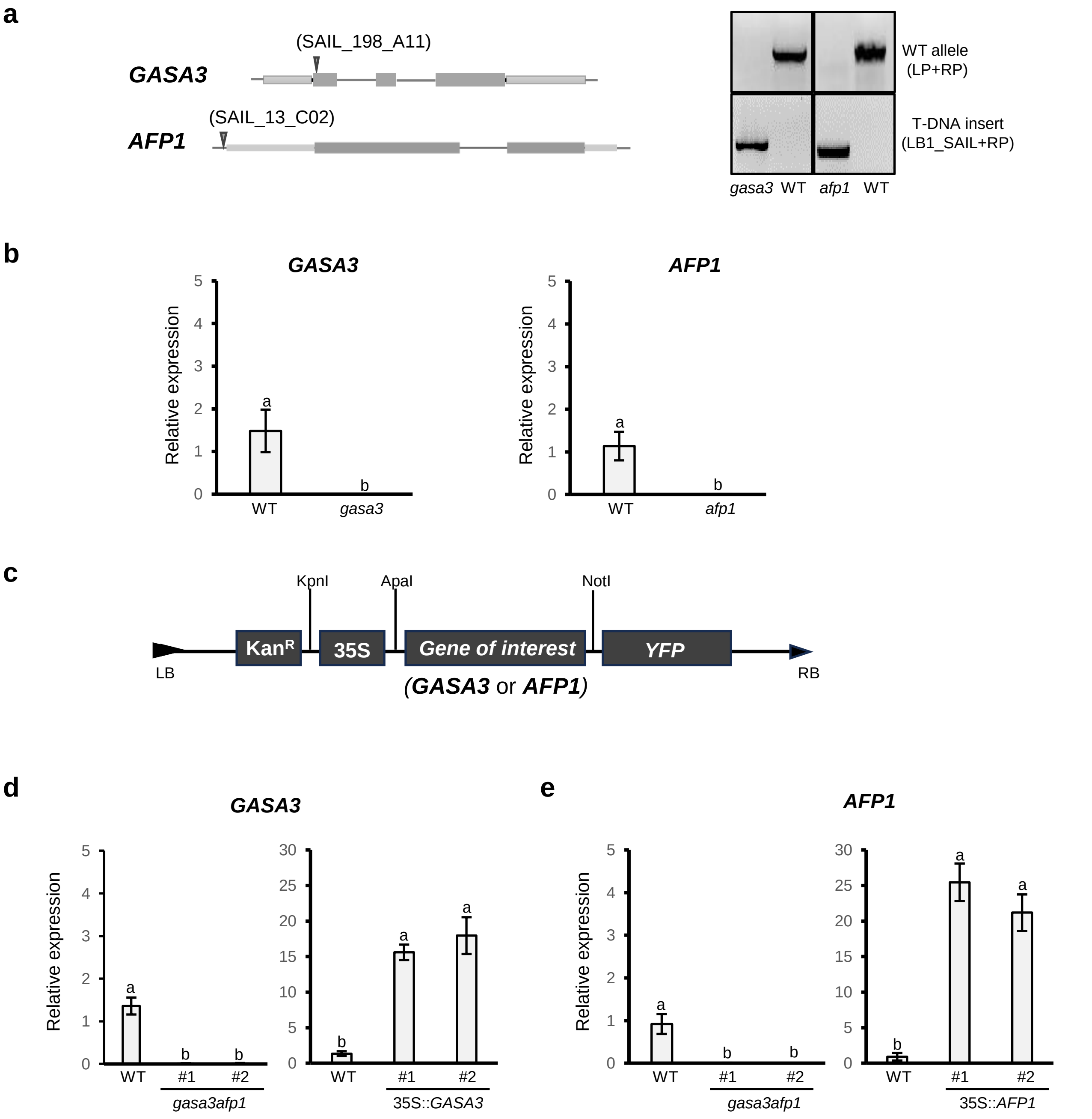

**Supplementary Figure S1: Characterisation of transgenic lines of *GASA3* and *AFP1*.**
(**a**) Schematic representation of the T-DNA insertion position in the lines *gasa3* (SAIL_198_A11) and *afp1* (SAIL_13_C02) (left panel). Confirmation of homozygous T-DNA insertion lines through PCR (right panel). (b) RT-qPCR confirmation of the *gasa3* and *afp1* null mutants. (c) Schematic representation of the p35S::(*GASA3 or AFP1)*-YFP cassette in pBIN19 vector used to produce overexpression and complementation lines. Transcript levels of (d) *GASA3* and (e) *AFP1* in *gasa3afp1* double mutants and p35S::*GASA3*-YFP and p35S::*AFP1*-YFP lines. For each two independent lines (#1 and #2) were tested. For RT-qPCR, RNA was harvested from rosettes of 32-day old Arabidopsis grown on soil under control conditions. Data represent means ± SE of three biological replicates (n=3), with statistical analyses carried out through one-way ANOVA and Tukey's Post-Hoc HSD tests (P<0.05).

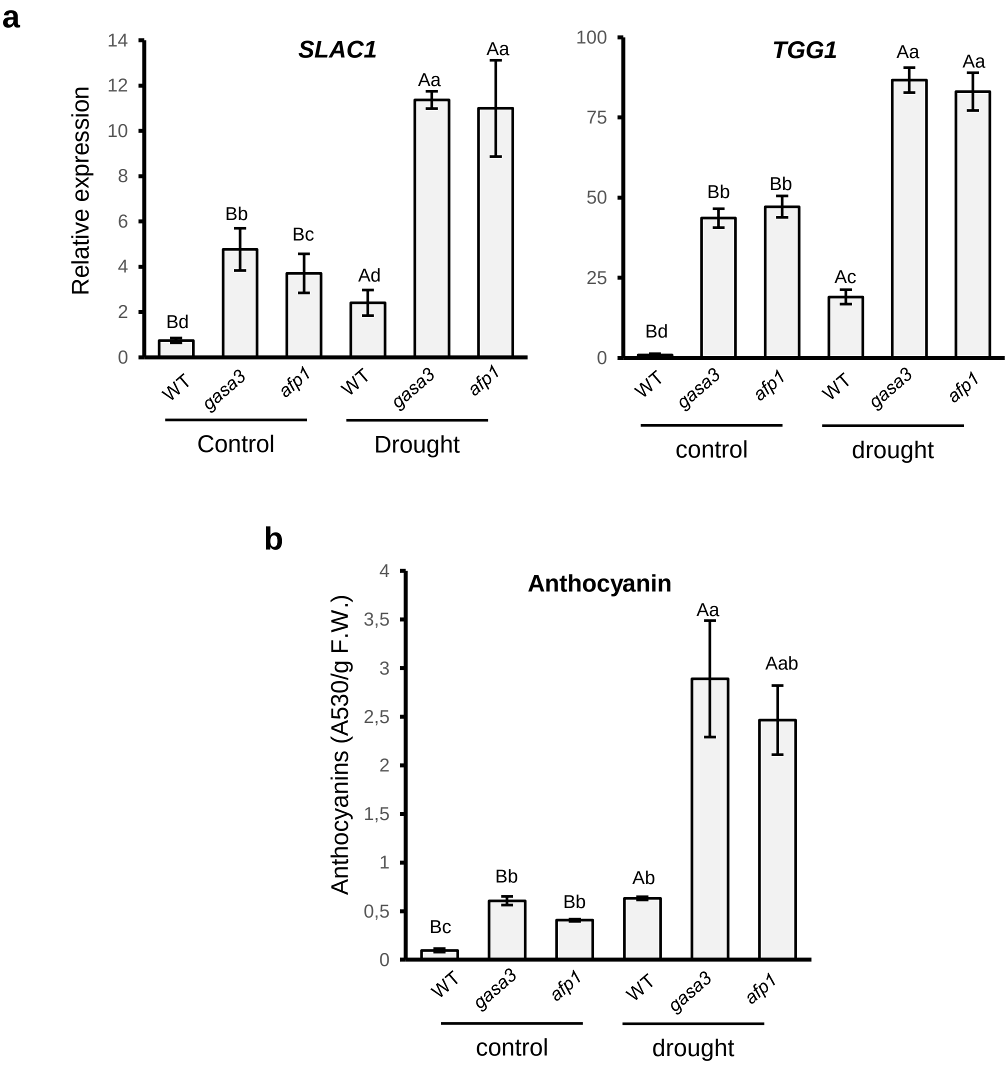

**Supplementary Figure S2: Effect of *GASA3* and *AFP1* on expression of genes related to drought tolerance.**

(a) Relative expression of *SLAC1* and *TGG1* as well as (b) anthocyanin contents of rosette leaves from WT, *gasa3* and *afp1* plants. All data shown were obtained from 32 day-old plants grown under control or progressive drought conditions (14 days of drought). Data represent means ± SE of three independent biological repeats (n=3). Statistical analyses were carried out with ANOVA and Tukey's Post-Hoc HSD tests ($P<0.05$).

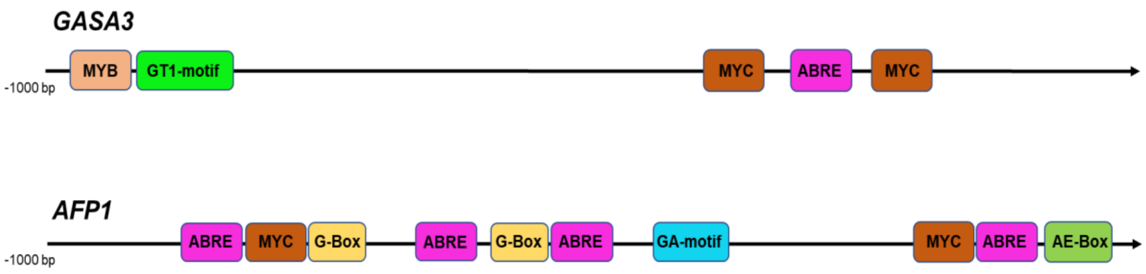

**Supplementary Figure S3 : cis-regulatory elementsfor abiotic stress in the promoter of *GASA3* and *AFP1*.**

Promoter analyses of *GASA3* and *AFP1* regions approximately 1kb upstream to the transcription start site was performed using Plant CARE (http://bioinformatics.psb.ugent.be). ABRE: ABA-responsive element; GT1-motif, G-box, GA-motif and AE-box: light-responsive elements. MYC- and MYB: elements involved in drought and ABA response.

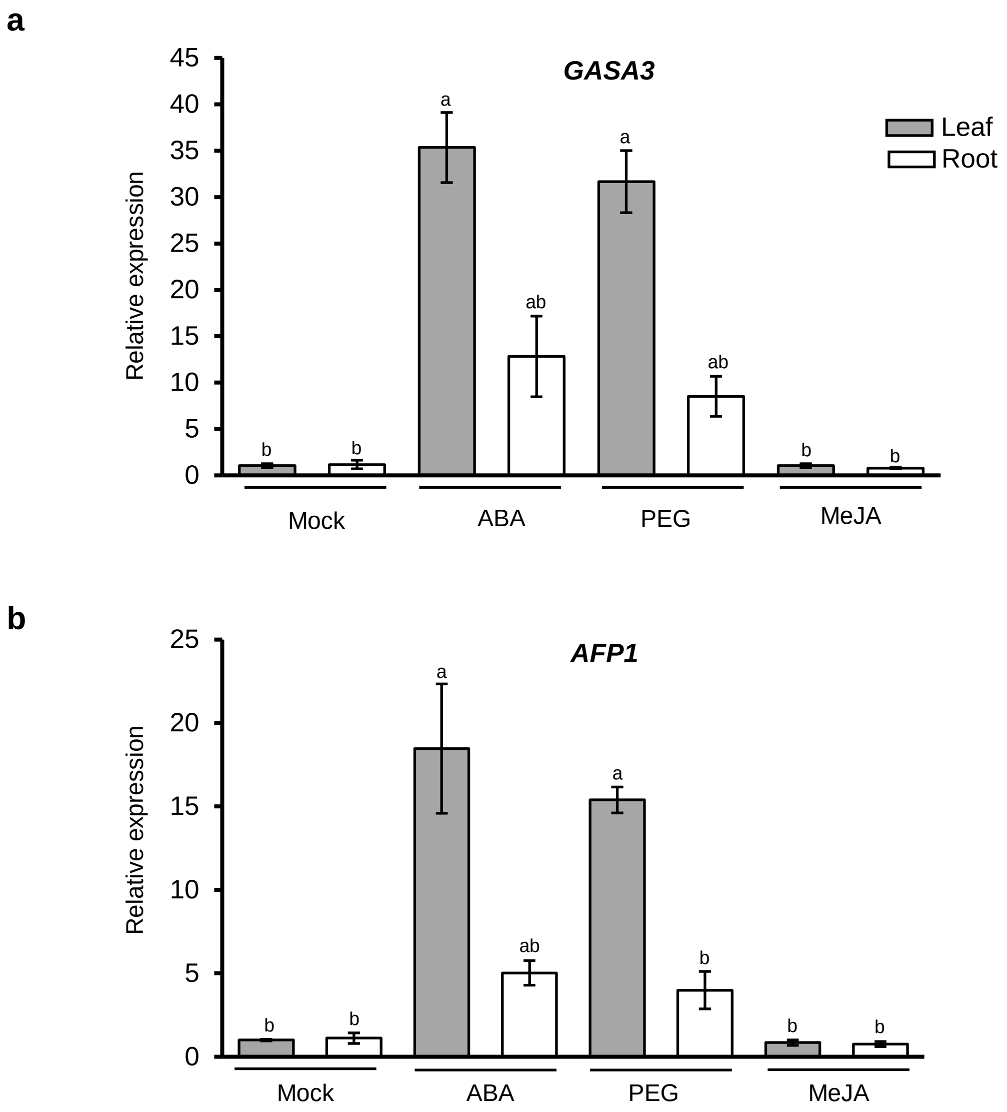

**Supplementary Figure S4: Regulation of *GASA3* and *AFP1* expression in leaves and roots.**
Relative expression of (a) *GASA3* and (b) *AFP1* in leaves and roots of 21-day old WT seedlings grown on ½ MS plates treated with either dd$H_2O$, 100 µM ABA, 100 µM MeJA, 100 µM $GA_3$ or 20% PEG-6000. Data represent means ± SE of three independent biological replicates (n=3). Statistical analyses were perfomed with one-way ANOVA and Tukey's Post-Hoc HSD tests ($P<0.05$).
